# Methanol-specific methyltransferase isozymes have large carbon kinetic isotope effects that impact methane isotopic signatures

**DOI:** 10.64898/2026.04.02.716163

**Authors:** Jonathan Gropp, Daniel A. Stolper, Dipti D. Nayak

## Abstract

The stable hydrogen and carbon isotopic composition of methane is widely used to determine its sources. Methanogenic growth on methanol generates methane with significantly lower ^13^C/^12^C ratios relative to other substrates, which is often used as a marker for this metabolism in environmental samples. The biochemical basis for the unusual isotope effect associated with methanol growth is currently unknown. Here, we grew *Methanosarcina acetivorans* on methanol and measured the change in the carbon and hydrogen stable isotopic compositions of the methane. We coupled these results with an inverse modeling approach to calculate the kinetic isotopic effects (KIEs) of the rate-limiting step, catalyzed by the methanol-specific methyltransferase complex (MTA). Through this process, we estimate the carbon KIE of MTA (^13^ε_MTA_) as −65.5 ‰ and the hydrogen KIE of MTA (^2^ε_MTA_) as −56 ‰. Next, we show that the ^13^ε_MTA_ contributes substantially to the large isotopic effect observed for methylotrophic methanogenesis on methanol. We also show that mutant strains that express only a single copy of the MTA complex (either MtaC1B1A1, MtaC2B2A1, or MtaC3B3A1) have ^13^ε_MTA_ and ^2^ε_MTA_ that are indistinguishable from the wild-type strain. Finally, based on a thermodynamic analysis, we propose that methanol activation by MTA will remain rate-limiting, even at low environmental methanol concentrations, and the large ^13^ε_MTA_ would be expressed in situ as well.

**Importance:** Methane is a potent greenhouse gas, and distinguishing between its biological sources is vital for modeling global carbon cycles. Methylotrophic methanogenesis produces methane with a uniquely depleted carbon isotope signature. However, the biochemical mechanisms driving this fractionation have remained unclear. In this study, we identify the methanol-specific methyltransferase (MTA) complex as the primary driver of these large carbon isotope effects. By utilizing *Methanosarcina acetivorans* mutants, we demonstrate that these effects are consistent across different MTA isozymes. Our results suggest these signatures are intrinsic to the enzyme complex and persist at low substrate concentrations. These findings provide a critical biochemical foundation for using stable isotopes to track microbial methane production in diverse natural ecosystems.

## Introduction

The stable isotopic ratios of carbon (^13^C/^12^C) and hydrogen (^2^H/^1^H) are used to trace biological and geological sources of methane in the environment (e.g., 1–3). These isotopic ratios are depicted using the δ notation:

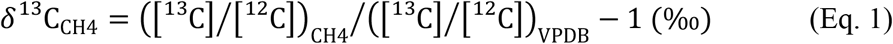

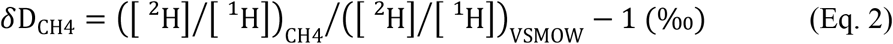

The δ values are reported relative to the VPDB (Vienna Pee Dee Belemnite) and VSMOW (Vienna Standard Mean Ocean Water) isotopic reference scales for carbon and hydrogen, respectively. Methanogenic archaea (methanogens) are the primary source of biogenic methane, and the δ^13^C_CH4_ and δD_CH4_ of the methane that they produce can vary substantially (1). This variability stems, in part, from the enzyme-specific kinetic isotope effects (KIEs) associated with catabolic enzymes involved in methanogenesis, that can vary in a substrate-dependent manner (4). These KIEs are defined as the ratio of the rate constants for a given reaction for the heavy vs. light isotope (^*h*^k and ^*l*^k, respectively, where KIE = ^*h*^*k*/ ^*l*^*k*). In addition to the KIEs, the δ^13^C_CH4_ and δD_CH4_ also depend on the pathway topology, i.e., if there are any branchpoints where carbon is diverted to an end product other than methane, and the degree of reversibility for a given enzyme, which is set by the free energy change associated with the reaction catalyzed (ΔG) (4–7). The extent to which each of these parameters influences δ^13^C_CH4_ and δD_CH4_ depends on factors such as substrate availability, temperature, and pH (6, 8–10). Under substrate-replete conditions, such as those typically used for laboratory batch cultures, enzymatic KIEs and pathway topology are the main determinants of δ^13^C_CH4_ and δD_CH4_ (7). To isolate the effect of KIEs and pathway topology from the δ^13^C of the substrate (δ^13^C_substrate_) under laboratory conditions, we can measure the isotopic difference between the substrate and methane using the *ε* notation:

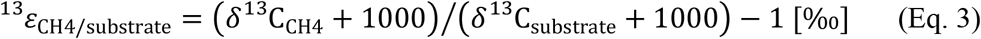

In laboratory batch cultures, methanogenic growth on methanol produces methane with the largest measured ^13^ε_CH4-substrate_ values (between ≈−65 ‰ to ≈−85 ‰) (4, 11–13), compared to other substrates that produce methane with a ^13^ε_CH4-substrate_ ranging between ≈−35 ‰ to ≈−70 ‰ (Fig. 1A) (4, 6, 8, 12, 14–26). Methanogenic growth on methanol is the major source of methane in environments such as salt marshes and mangrove forests (e.g., 27–34), and the distinct ^13^ε_CH4-substrate_ values observed for this pathway could be critical to assess its contribution to global methane budgets. However, the molecular and biochemical rationale for these large isotopic offsets are poorly understood and are the focus of this study.

**Fig. 1.**
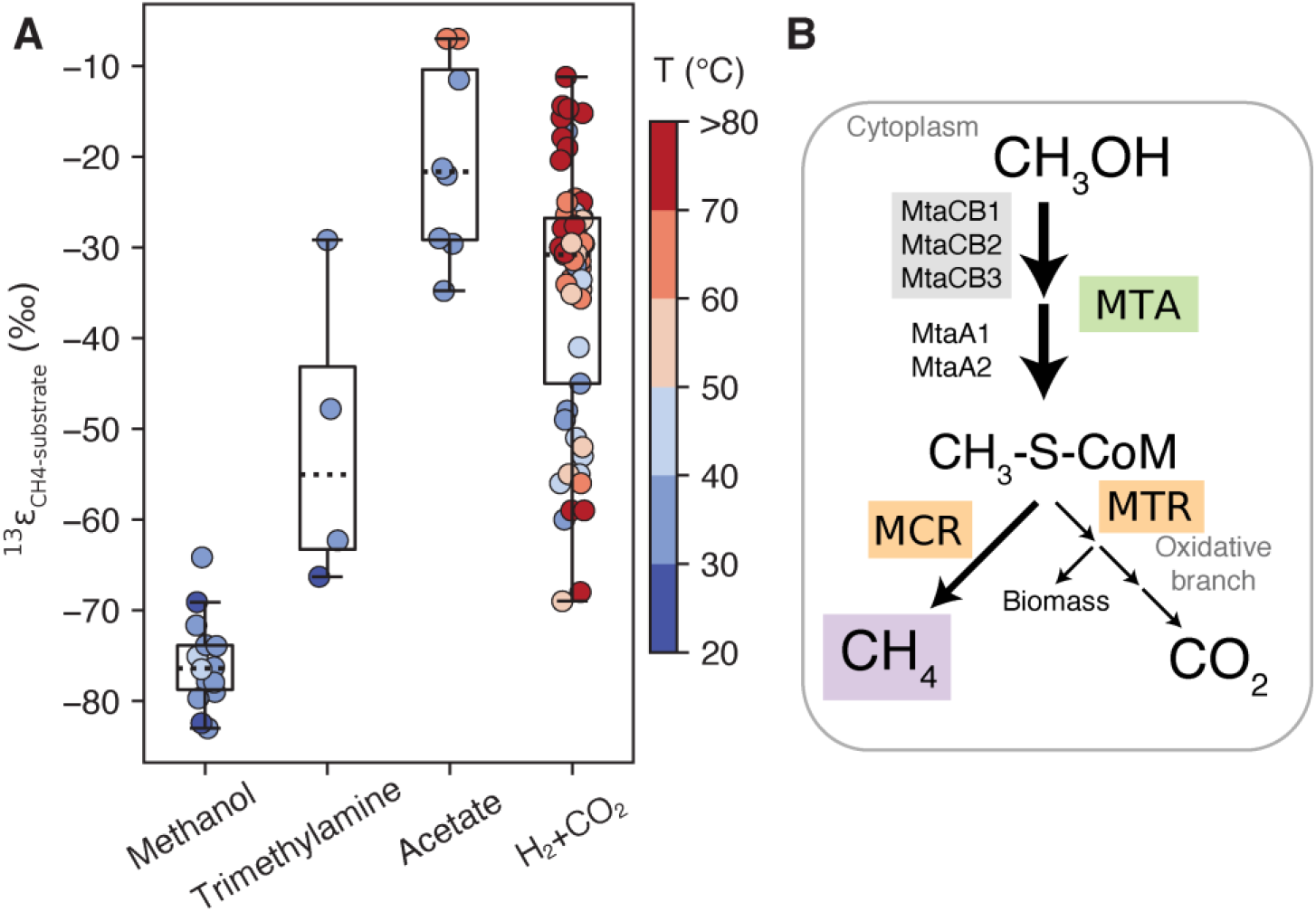
Isotopic effects of microbial methane production. **(A)** The ^13^ε_CH4-substrate_ for different methanogenic carbon sources in axenic laboratory cultures. The box plots represent the 25^th^, 50^th^, and 75^th^ percentiles of the data. The colors represent the growth temperature (Data from Refs. (4, 6, 8, 12, 14–26), see full list in Table S1). We do not provide data for hydrogen isotopes as the hydrogen isotopic compositions of methane generated from methylated compounds and acetate is controlled by the δD of both the organic substrate and the water. Hence the hydrogen isotopes can vary depending on the δD of the water used for growth. **(B)** Our simplified metabolic pathway for methylotrophic methanogenesis on methanol in *Methanosarcina acetivorans*. The methyl group from methanol is activated by a methanol-specific methyl-transferase complex (MTA) to methyl-coenzyme M (CH_3_-S-CoM). The complex includes MtaCB, which has three isoforms in *M. acetivorans*, and MtaA, which has two isoforms. The CH_3_-S-CoM is either reduced to CH_4_ or oxidized to CO_2_ in a 3:1 ratio. The more detailed metabolic pathway used for data fitting is given in Fig. S1. Abbreviations for enzyme complexes names: MTR – N^5^-methyltetrahydromethanopterin:coenzyme M methyltransferase; MCR – methyl-coenzyme M reductase.

During methanogenic growth on methanol, the substrate likely diffuses across the cell envelope to enter the cytosol, where a methanol-specific methyltransferase (MtaB) catalyzes a methyl-transfer reaction to a corrinoid-containing protein, MtaC (Fig. 1B). There are three copies of the MtaCB complex in the model organism used for this study, *Methanosarcina acetivorans*, which are encoded by *mtaC1B1, mtaC2B2*, and *mtaC3B3* (Fig. 1). All three MtaCB enzymes in *M. acetivorans* catalyze the same biochemical reaction, i.e., they are isozymes, and are expressed during methanol growth under laboratory conditions (35, 36). The methyl-group bound to MtaC is subsequently transferred to coenzyme M (CoM-SH) to form methyl–coenzyme M (CH_3_-SCoM) by a second methyltransferase (MtaA) (Fig. 1B). There are two copies of MtaA in *M. acetivorans*, which are encoded by *mtaA1* and *mtaA2* (Fig. 1). Because the two reactions act in concert (Fig. 1B), here we refer to the combination of MtaCB and MtaA as the methyltransferase complex (MTA) in this work. Without any biomass production or any other products, the methyl groups in CH_3_-S-CoM must produce methane and carbon dioxide in a 3:1 ratio to maintain redox balance. The reduction of CH_3_-S-CoM to methane is catalyzed by MCR, and the stepwise oxidation of the methyl group in CH_3_-S-CoM to CO_2_ is catalyzed by a series of six enzymes that we collectively term the “oxidative branch” (Fig. 1B and Fig. S1).

MTA is the only enzyme complex in the methanogenic pathway that is unique to methanol metabolism and mediates the first enzymatically controlled step in this process. Based on this, Krzycki et al. (4) proposed that the KIE of MTA underlies the large ^13^ε_CH4-substrate_ values observed on methanol. Penger et al. (13) also considered this hypothesis but proposed an additional hypothesis wherein the disproportionation of CH_3_-S-CoM to CH_4_ and CO_2_ contributes to the large ^13^ε_CH4-substrate_ values associated with methanogenesis on methanol. However, this branched pathway topology is a common feature of methylotrophic methanogenesis (methanogenic growth on methylated substrates) on other substrates like trimethylamine that do not produce such large ^13^ε_CH4-substrate_ values (Fig. 1A). Taken together, we consider it unlikely that the branched topology of methylotrophic methanogenesis is the origin of the large ^13^ε_CH4-substrate_ observed for growth on methanol (13). Rather, we propose that the more parsimonious explanation is that MTA is the isotopically discriminating step in the pathway under typical laboratory growth conditions. It is important to note that if MTA controls ^13^ε_CH4-substrate_, then the isotopic composition of methane is set by a composite of the three isozyme-specific isotopic effects.

To estimate the KIE of MTA, we built a quantitative isotopic model of methylotrophic methanogenesis by placing constraints on the KIEs of key enzymatically catalyzed steps in the pathway. At present, experimentally derived KIEs are available only for one enzyme, methylcoenzyme M reductase (MCR) (37), which catalyzes methane production in the final step of methanogenesis. Rather than purify each of the remaining enzymes and measure their KIEs in vitro (e.g., 37), here we employ an inverse modeling approach to determine the KIEs with the assumption that, under substrate-replete conditions, i.e., under limited reversibility, there is a single rate-limiting step for methanol-mediated methanogenesis (38). We then employ the same approach to mutants of *M. acetivorans* encoding either *mtaC1B1A1, mtaC2B2A1*, or *mtaC3B3A1* to estimate the KIEs of each MtaCB isozyme and compare these KIEs to the wild-type strain which encodes all three isozymes. We find that the calculated carbon KIEs for each MtaCB isozyme are relatively large (−65‰) and similar across the isozymes. Overall, our findings provide experimental support for the hypothesis that the large carbon KIEs of MTA dominate the carbon isotope effects observed during methanogenic growth on methanol.

## Results

### Growth parameters and isotopic compositions of methane from *M. acetivorans* strains that express single methyltransferase isozymes

All growth experiments were conducted at 37 °C in batch culture in hermetically sealed Balch tubes containing 10 mL minimal high-salt (HS) medium supplemented with 125 mM methanol as the sole carbon and energy substrate. We designate WWM1 (39) as the “wildtype” strain of *M. acetivorans* (WT), as it is the parent for mutant strains expressing only one of the three *mtaCB* operons under a minimal *mcrB* promoter (WWM170, WWM176, WWM184; Ref. (35), genotypes listed in Table 1). All three mutant strains had ∼2-fold slower growth rates and ∼5-fold longer lag times relative to the WT, as shown previously (Fig. 2A-E and Tables S2 and S3) (35).

**Table 1.**
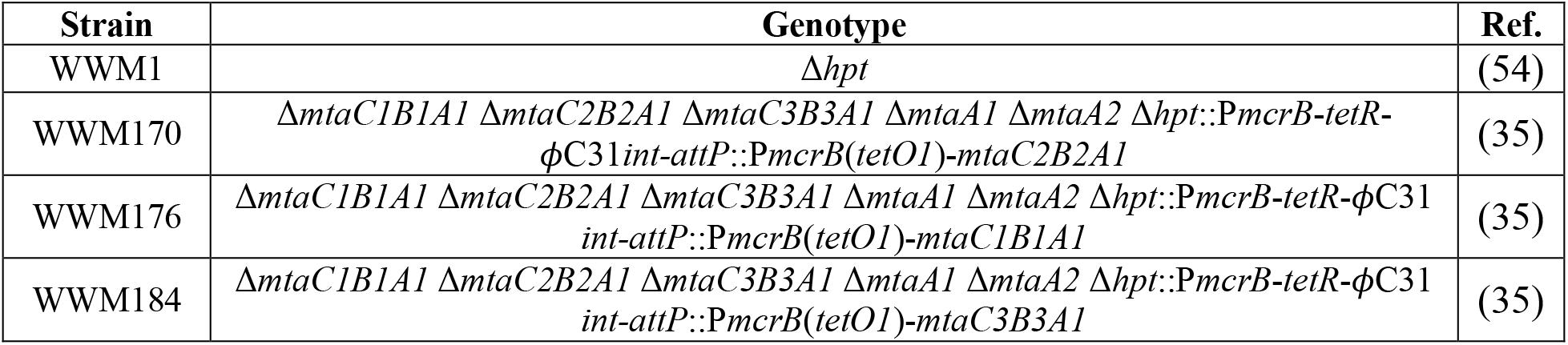
*Methanosarcina acetivorans* strains used in this study.

| Strain | Genotype | Ref. |
| --- | --- | --- |
| WWM1 | $\Delta hpt$ | (54) |
| WWM170 | $\Delta mtaC1B1A1 \Delta mtaC2B2A1 \Delta mtaC3B3A1 \Delta mtaA1 \Delta mtaA2 \Delta hpt::PmcrB-tetR-\phi C31int-attP::PmcrB(tetO1)-mtaC2B2A1$ | (35) |
| WWM176 | $\Delta mtaC1B1A1 \Delta mtaC2B2A1 \Delta mtaC3B3A1 \Delta mtaA1 \Delta mtaA2 \Delta hpt::PmcrB-tetR-\phi C31int-attP::PmcrB(tetO1)-mtaC1B1A1$ | (35) |
| WWM184 | $\Delta mtaC1B1A1 \Delta mtaC2B2A1 \Delta mtaC3B3A1 \Delta mtaA1 \Delta mtaA2 \Delta hpt::PmcrB-tetR-\phi C31int-attP::PmcrB(tetO1)-mtaC3B3A1$ | (35) |

**Fig. 2.**
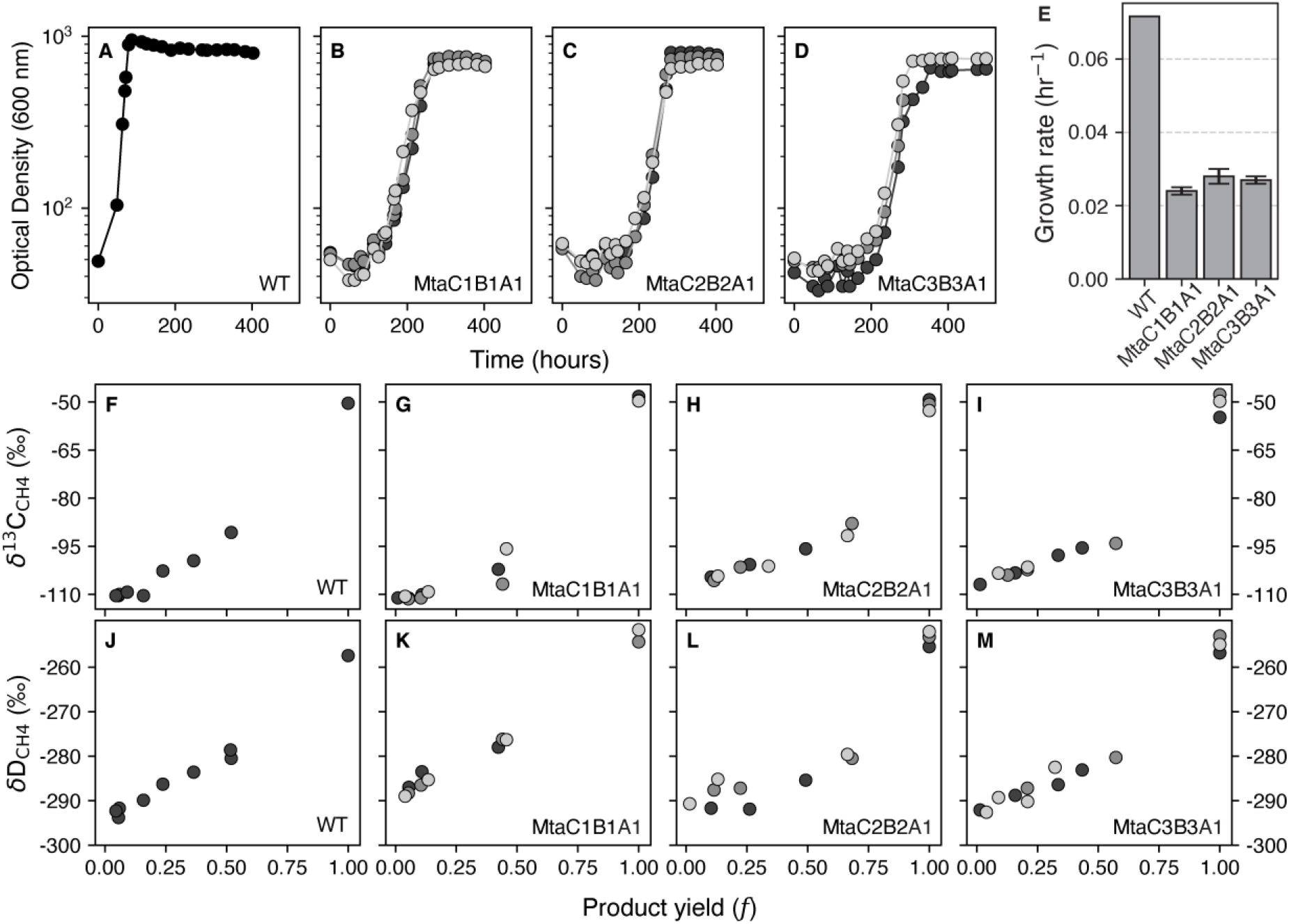
Growth, methane production, and isotopic data for *M. acetivorans* strains expressing different methyltransferase isozymes. (**A-D**) Representative growth curves where each color represents a single biological replicate. Cells were grown on high-salt (HS) minimal medium with 125 mM methanol at 37 °C. All growth curves are provided in Fig. S2. **(E)** Growth rates of WWM1 (WT), WWM176 (expressing only MtaC1B1A1), WWM170 (expressing only MtaC2B2A1), and WWM184 (expressing only MtaC3B3A1), where errors bars represent 1 standard error of the mean (SEM, *n*=3) (See Table 1 for list of strains used here). (**F-I**) δ^13^C_CH4_ and (**F-I**) δD_CH4_ values vs. the relative amount of CH_4_ produced (*f*). Each data point in **F-M** is a sacrificed culture (see details in Materials and Methods); the colors represent the replicate experiments. The initial measured δ^13^C and δD of the methanol were −41.6‰ and −49‰, respectively, and the measured δD of the water was −79‰.

We tracked the δ^13^C_CH4_ over the course of a growth cycle using isotope-ratio mass spectrometry (IRMS). The δ^13^C_CH4_ increased as methane production approached completion, i.e., *f = 1*, as expected for Rayleigh isotopic fractionation in a closed system. The δ^13^C_CH4_ at stationary phase, i.e., when the methanol was fully consumed, was comparable between WT and the mutants, and the values ranged between −55‰ and −48‰. These values are lower than the δ^13^C of the methanol used for these growth experiments (−41.7‰) (Fig. 2F-I). This difference in the δ^13^C is within the range observed in previous studies with *M. acetivorans* as well as with other methanogens grown on methanol (δ^13^C_CH4_ 8‰–12‰ more negative than δ^13^C initial methanol) (9, 13, 25, 40), and is typical for a branched metabolic pathway where the substrate is converted to multiple products that have different isotopic compositions (5).

Like δ^13^C_CH4_, the hydrogen isotopic composition of methane (δD_CH4_) increased with methane production. At stationary phase, the δD_CH4_ was also similar between WT and the mutants too, and the values ranged between −257‰ and −252‰ (Fig. 2J-M). The final δD_CH4_ at stationary phase is set by both the isotopic compositions of the water in the growth medium, which in our experiments had a δD of −79‰, and the initial δD of the methyl group of the methanol, which had a δD of −49‰ (see Materials and Methods). Consequently, and unlike the carbon isotope data, the δD_CH4_ cannot be compared across studies in a straightforward manner as the δD_H2O_ of the growth medium in different laboratories can vary substantially (range of ≈160‰, 41)

### A Bayesian approach to estimate KIEs in a physiological context

We calculated enzymatic KIEs using the measured δ^13^C_CH4_ and δD_CH4_ values (Fig. 2) in the context of a published reaction network of methylotrophic methanogenesis (9). This reaction network includes the activation of methanol catalyzed by MTA to generate CH_3_-S-CoM, its subsequent disproportionation to methane and CO_2_, as well as a biomass sink of 10% (Fig. 1B and Fig. S1). We assume a constant ratio of 3:1 between CH_4_ and CO_2_ over the course of a growth cycle. Based on a previously published metabolic model of the closely related strains *Methanosarcina barkeri*, we assume that the MTA- and MCR-catalyzed reactions are irreversible and that methanol diffusion is sufficiently rapid that it neither limits growth nor sets final methane isotopic compositions (38). We permit a partial reversibility of 0.05 in the oxidative branch based on a prior study where we focused on constraining degrees of reversibility of the pathway using the same growth conditions as used here (9). We verified that this choice did not affect final conclusions via sensitivity testing (see below). We also tested the sensitivity of the metabolic model to the chosen degree of reversibility in MCR and in MTA.

The metabolic model employed in this study can be described mathematically as a series of ordinary differential equations that quantify the rates of production and/or consumption of CH_3_OH, methane, and CO_2_ (see Materials and Methods) and the respective mass balance for the isotopically unsubstituted and singly substituted species. To run the model, the initial δ^13^C and δD values of the methanol (δ^13^C_CH3OH_ and δD_CH3OH_, respectively) and the δD of the water must be independently known and we assume that the concentration and isotopic compositions of the intermediates are at a steady state between integration steps. The latter is a common assumption of isotopic models (42). The output of the model includes the time-dependent concentrations for CH_3_OH, CO_2_, CH_4_, and their respective δ^13^C_CH3OH_, δ^13^C_CO2_, δ^13^C_CH4_, and δD_CH4_ values. As growth occurs in hermetically sealed tubes, we assume the system is closed such that all product methane accumulates and mixes in the system during growth.

We used our isotopically-enabled metabolic model to calculate the enzyme-specific KIEs for MTA using a Markov-Chain Monte-Carlo (MCMC) algorithm to fit the data as previously described (9) (see model parameters in Table S4). The algorithm generates posterior distributions of the KIEs that generate the best fit to the experimentally measured δ^13^C_CH4_ and δD_CH4_ values. Because KIEs are often small with values close to unity, we report them using the ε notation where *ε* = 1000 × (KIE − 1) (‰). To determine the carbon and hydrogen KIEs of MTA (^13^ε_MTA_ and ^2^ε_MTA_, respectively), we first defined the allowable boundaries for the KIEs, based on typical measured ranges in the literature (45). For carbon KIEs, we bound ^13^ε to be between −100‰ and 0‰ (0.9 < KIE < 1). For hydrogen KIEs of reactions that break or create a bond with the hydrogen atom (primary KIEs) we used a ^2^ε range between −900‰ and 0‰ (0.1 < KIE < 1). For hydrogen KIEs of reactions that do not break or create a bond with the hydrogen atom (secondary KIEs), we used a smaller range of ^2^ε between −300‰ and +200‰ (0.7 < KIE < 1.2). For these secondary isotope effects, we allow for the possibility of inverse effects in which deuterated species react faster than isotopically unsubstituted species. Using these boundary conditions, we then ran the MCMC algorithm with 20,000 total walker-steps for each experimental dataset (a total of 10 experiments, one for the WT and three for each mutant strain). We verified that our chosen bounds do not affect the best-fit values of the isotope effects (Fig. S3 and Fig. S4).

In addition to the KIEs of MTA, we have also calculated the ratio of the KIEs of MCR and MTR (^13^KIE_*MCR*_⁄^13^KIE_*MTR*_ ≡ ^13^KIE_*MCR*/*MTR*_), which controls the δ^13^C_CH4_ values at the end of the experiment when *f* = 1. Although our metabolic framework in principle allows us to constrain the KIEs of all reactions in the methanogenic pathway, we chose not to extend our analysis beyond MTA and the ^13^KIE_MCR/MTR_ because measurement of δ^13^C_CH4_ and δD_CH4_ values alone cannot not provide sufficient constraints to solve for additional free parameters downstream of these reactions.

### Large carbon kinetic isotopic effects associated with methanol-specific methyltransferases

Using the MCMC fitting algorithm outlined above, we calculated the ^13^ε_MTA_ values for WT and the three mutants expressing a single copy of each the three *mtaCB* operons (WWM170, WWM176, WWM184). The median ^13^ε_MTA_ for WT was 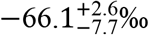, where the uncertainty represents the lower and upper bounds of 95% of the model results (which need not be symmetric). The median ^13^ε_MTA_ of the mutants that encode only a single *mtaCB* isoform was 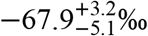 for MtaC1B1A1, 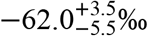 for MtaC2B2A1, and 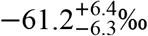 for MtaC3B3A1 (Table 2 and Fig. 3A–D). These are statistically indistinguishable from each other and WT (*p*-value > 0.05%, see Materials and Methods). We also used the MCMC fitting algorithm to analyze a previously published dataset for methanol growth at 37 °C for *M. acetivorans* (13), which yielded ^13^ε_MTA_ values of 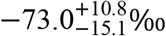 and 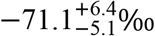, based on separately fitting with their measured δ^13^C_CH3OH_ and δ^13^C_CH4_ values (Table S5). Both values are also within the uncertainty range of our estimated ^13^ε_MTA_ for the WT and the mutant encoding MtaC1B1A1.

**Table 2.**
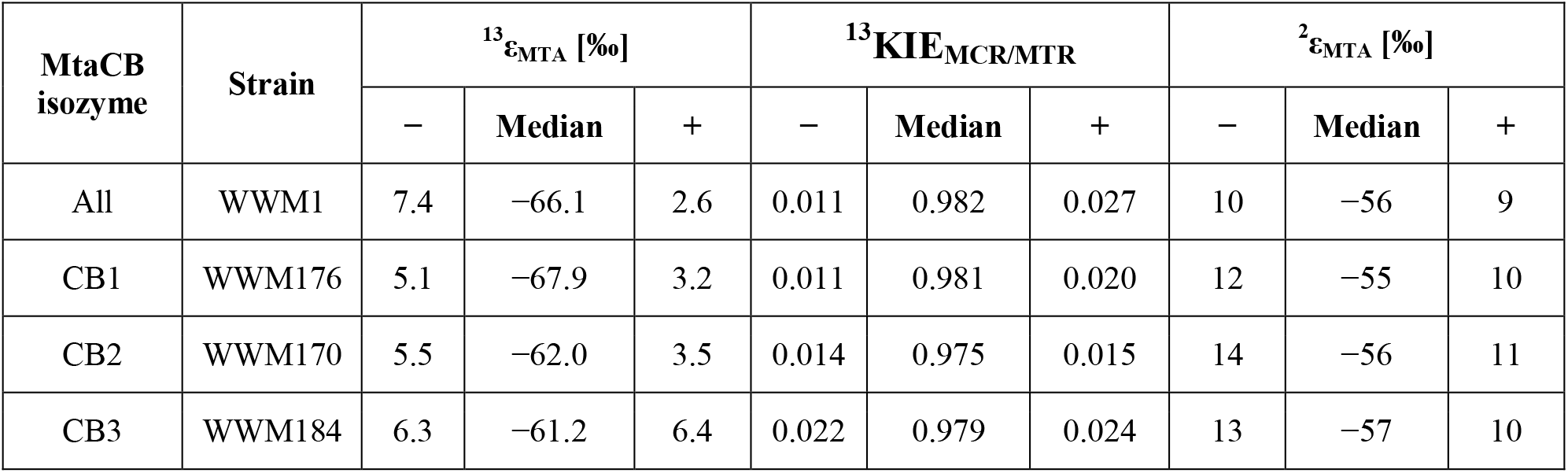
Calculated carbon and hydrogen KIEs for *M. acetivorans*. The values for the mutant strains (WWM170, WWM176, and WWM176) represent pooled posterior distributions of three replicate experiments, and the uncertainties represent 95% of the model results. The data for each replicate is shown in Table S6. The values of ^13^ε_MTA_, ^2^ε_MTA_, and ^13^KIE_MCR/MTR_ were statistically indistinguishable between the strains (see Materials and Methods).

**Fig. 3.**
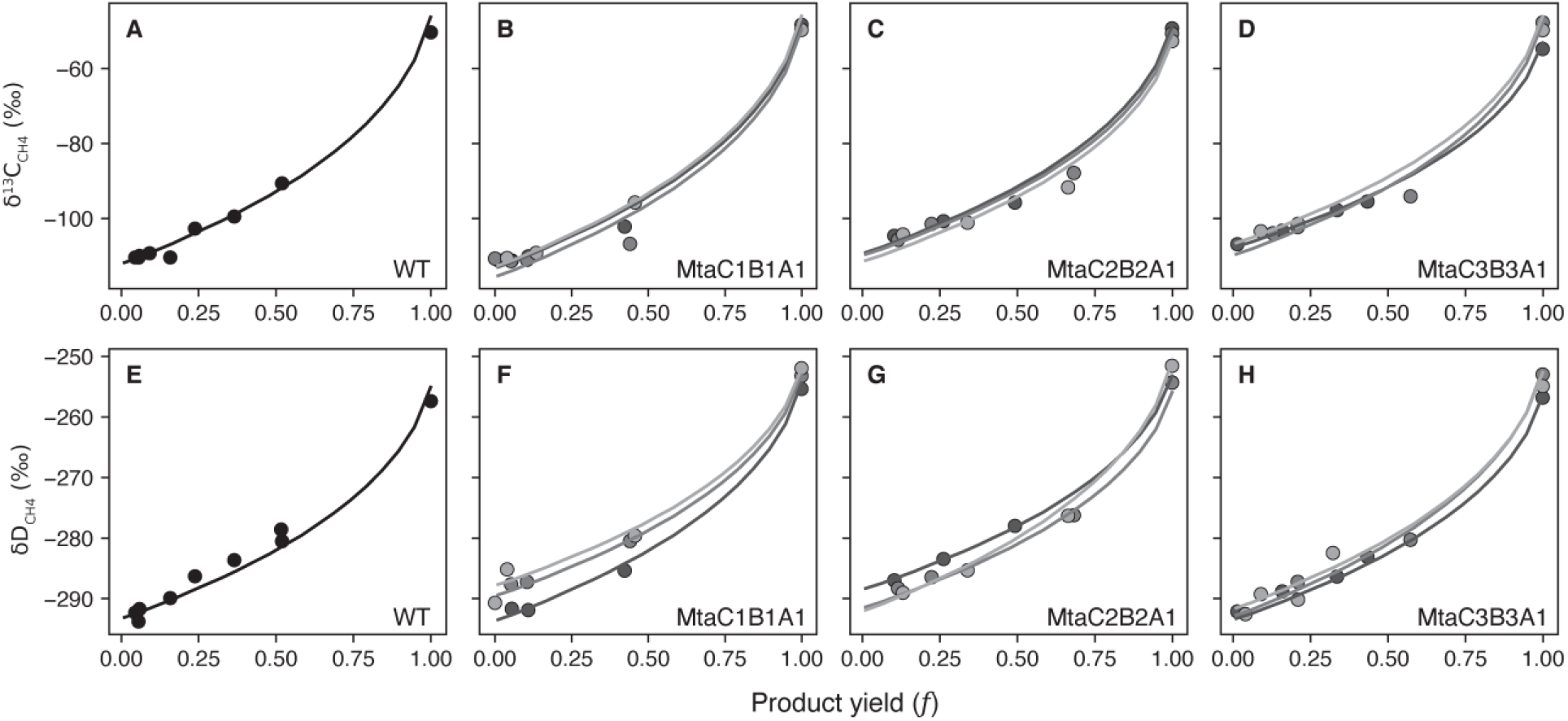
Isotopic composition of methane in the wildtype and mutants that express single methanol-dependent methyltransferase isozymes. Measured (circles) and model fit (lines) (**A-D**) δ^13^C_CH4_ and (**E-H**) δD_CH4_ vs. methane production yield. Symbols represent methane measured in the headspace of a single tube, and the gray shades represent replicates of the mutant strains. The model-fitted δ^13^C_CH4_ and δD_CH4_ values were obtained using the median values of the posterior KIEs as given in Table 2.

Unlike MTA, we cannot estimate the absolute values of the carbon KIEs of MCR and MTR. To do so requires additional experimental data, like the isotopic composition of labile metabolic intermediates (like CH_3_-S-CoM) in the methanogenic pathway, which is not currently possible. However, the model does allow us to estimate the ratio of the KIEs of the two enzymes (^13^KIE_MCR/MTR_). The ^13^KIE_MCR/MTR_ of 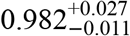 for the WT is indistinguishable from the mutant strains with ^13^KIE_MCR/MTR_ of 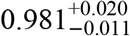 for MtaC1B1A1, 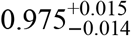 for MtaC2B2A1, and 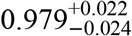 for MtaC3B3A1 (Table 2). If we use the previously measured value of ^13^ε_MCR_ ≈−40±10‰ from *Methanothermobacter marburgensis* (37), we obtain ^13^ε_MTR_ values between −15‰ to −22‰ for ^13^KIE_MCR/MTR_ values of 0.975 and 0.982, respectively, which are the lowest and highest median values that we calculated. Since *M. marburgensis* is a thermophilic methanogen that grows at 60 °C, it is not currently known if the ^13^ε_MCR_ from this strain is representative for a distantly related, mesophilic methanogen like *M. acetivorans*. To constrain ^13^ε_MTR_ accurately, we would require an independent estimate of ^13^ε_MCR_ of the enzyme purified from *M. acetivorans* at 37 °C, which is beyond the scope of this study.

We conducted a variety of sensitivity analyses to test whether various model assumptions impact the results presented above. First, we tested whether the size of the biomass sink and the degree of reversibility of the oxidative branch affect the ^13^ε_MTA_ and ^13^KIE_MCR/MTR_ values (Table S6). We found the choice of the size of the biomass sink had a negligible impact on the calculated ^13^ε_MTA_ values (<1.8‰) and on the ^13^KIE_MCR/MTR_ values (<0.009) (Table S6). Specifically, these changes are within the 95% uncertainty range of our model results and would have no impact the median values or the outcome of statistical tests. Similarly, changing the reversibility in the oxidative branch between 0 (i.e., irreversible) to 0.1 had a small effect on the ^13^ε_MTA_ (<1.9‰) and ^13^KIE_MCR/MTR_ values (<0.008), which were also within the uncertainty bounds of the model results. Relaxing the assumption that MTA is irreversible by increasing its reversibility to 0.2, changed the ^13^ε_MTA_ by a value up to 4‰, which was also within the uncertainty bounds of our model (Table S6).

### Small-normal hydrogen kinetic isotopic effects associated with methanol-specific methyltransferases

We used our MCMC algorithm to also calculate the hydrogen KIE of MTA (^2^ε_MTA_). This is a secondary isotope effect as the methyltransferase reaction does not break or create any new C-H bonds. Secondary hydrogen KIEs are typically smaller in absolute magnitude than primary isotope effects (43). Our model predicts that the ^2^ε_MTA_ is normal, with a value of 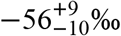 for the WT strain, which is indistinguishable from values for all mutant strains: 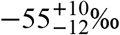 for MtaC1B1A1, 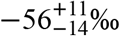 for MtaC2B2A1, and 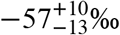 for MtaC3B3A1 (Table 2 and Fig. 3E–H). To the best of our knowledge, these are the first estimates of ^2^ε_MTA_ and so there are no other experiments to which we can compare our estimates.

Unlike carbon isotope effects, we cannot estimate the ratios of the hydrogen KIEs for MCR and MTR. This is because this ratio depends on three parameters: the KIE of MTR and the primary and secondary KIEs of MCR. As shown in Fig. S4, the dependence on KIEs does not allow for a unique solution of any of their ratios. Finally, we conducted a sensitivity test of our assumption that MTA is irreversible. Specifically, we calculated ^2^ε_MTA_ for MTA’s reversibility from 0 (irreversible) to 0.2 and found that increasing the reversibility of MTA lowers the KIE, i.e., makes ^2^ε_MTA_ less negative, by up to 34‰ (Table S6). While this is a significant change, it does not impact our overall conclusion that ^2^ε_MTA_ is a normal isotope effect that is small (^2^ε_MTA_ ∼ −60‰) in magnitude.

## Discussion

Methylotrophic methanogens growing on methanol have long been known to express large carbon isotope effects. The combination of our experimental data and isotope-enabled metabolic model indicate that these effects stem from the kinetically-controlled and irreversible uptake of methanol by MTA. Specifically, we find that the isozyme-specific ^13^ε_MTA_ values range between −61.2‰ to −67.9‰ and contribute to ≈90% of the overall carbon isotopes effect relative to the source methanol (Table S7), with the rest coming from the branching point between MCR and the oxidative pathway. These results support a prior hypothesis that large isotopic effects in methylotrophic methanogenesis emerge due the KIE of the substrate-specific methyltransferase reaction (4). By performing growth experiments and isotopic measurements on WT and mutants expressing each of the single MtaCB isozymes, we can also conclude that all of the isozymes have similar KIEs (Table 2). The δ^13^C_CH4_ values at the end of the growth experiments were consistently lower than the initial δ^13^C_CH3OH_ values by ≈9‰ (Table S2) and accordingly, we found that ^13^ε_MCR_ is consistently larger in absolute magnitude than ^13^ε_MTR_, with KIE ratios translating to a difference of ∼15‰. However, this difference contributed to only ≈10% of the overall carbon isotopic offset between methane and methanol (Table S7).

We first discuss the implications of the results for the biochemistry of the enzymes. Isotopic effects allow for the identification of reaction mechanisms (43, 44). The methylation of the MtaC-bound cob(I)amide catalyzed by MtaB is thought to proceed via an S_N_2 reaction mechanism, in which the carbon in the methanol is activated by Zn(II), a strong electrophile, in MtaB and then attacked by the strong nucleophile, a cob(I)amide in MtaC (45). S_N_2 methylation reactions commonly have small (in absolute terms) normal or inverse secondary hydrogen KIEs that range between −100‰ and +100‰ (46). This contrasts with S_N_1 mechanisms, which commonly have larger normal secondary hydrogen KIEs of up of −350‰ (46). In addition, S_N_2 reactions often have a relatively large carbon KIEs that range between −40‰ and −80‰ (46–49). Thus, our findings of a small hydrogen KIE (≈−55‰) and large carbon KIE (≈−65‰) support an S_N_2 reaction mechanism for the MtaB-mediated methyltransferase reaction.

We now turn to discussing the environmental implications of our estimated KIEs. Changes in environmental conditions impact isotopic composition of methane as has been clearly demonstrated for hydrogenotrophic methanogens (e.g., 8). Under high nutrient conditions in the laboratory, hydrogenotrophic methanogens generate methane with isotopic compositions that are similar to those detected in high energy environments (such as near-surface terrestrial systems), not low energy environments (e.g., deep-sea sediments) (50). These differences have been attributed to changes in enzyme reversibility as a function of the amount of free energy (ΔG) available in the environment to drive methanogenesis (7–9, 51).

It has also been proposed that methylotrophic growth at lower absolute ΔG in the environment can lead to the expression of equilibrium hydrogen isotope effects driven by enzyme reversibility (52). It is unclear if this is also the case also for carbon isotopes. The theoretically determined carbon isotopic equilibrium between CH_4_ and methanol is smaller than ^13^ε_methanemethanol_ observed in laboratory cultures, though with the same sign, with methane lower in ^13^C/^12^C vs. the substrate (∼−20‰ vs ∼−70‰, respectively. Ref. 48). Isotopic equilibration between substrate methanol and product methane requires that MTA is fully reversible. However, MTA is predicted to be <1% reversible, (i.e., effectively irreversible) down to environmentally-relevant methanol concentrations (∼1 μM, 38). For context, high-sulfate marine sediments in the Gulf of Mexico have methanol concentrations between 1 and 10 μM and are dominated by methylotrophic methanogenesis. In such environments, δ^13^C_CH4_ values as low as ≈−90‰ are observed vs. a δ^13^C of ≈−20‰ for the total organic carbon, which we use as the approximate maximum value for methanol generated from fermentation of the organic carbon (27). As such, we propose it unlikely that high degrees of reversibility in MTA can drive equilibration of either carbon or hydrogen isotopes.

Importantly, it is possible to modify the isotopic composition of methane without altering the reversibility of MTA. This can be done by increasing reversibility in the oxidative branch of the metabolic pathway or in MCR, either of which could result in hydrogen and carbon isotope exchange and even equilibration between methane and water for hydrogen isotopes and methane and CO_2_ for carbon isotopes, regardless of the reversibility of the MTA-catalyzed reaction (8, 9). Reversibility in the branched portion of the network could occur given a sufficiently low environmental ΔG, for example due to low substrate concentrations. We propose that such equilibration is more likely for hydrogen isotopes and less so for the carbon isotopes. This is because there are four different enzymes that exchange hydrogen isotopes with water and just two of them would overwrite the KIEs of MTA if the reversibility was high enough. In contrast, exchange of carbon isotopes would require all the steps between CH_4_ and CO_2_ to be reversible (Fig. S1), which is possible but requires lower ΔG gradients (i.e., closer to 0) (7, 8). This decoupling between hydrogen and carbon isotopes has implications for interpreting δD_CH4_ and δ^13^C_CH4_ values of methylotrophic methanogenesis in natural environments. Specifically, we predict that methylotrophic methanogenesis from methanol will have a large carbon isotope offset across a wide range of growth rates while the δD_CH4_ values would trend towards isotopic equilibrium with the water as the free energy available to the cells decreases (9).

Finally, we have shown here the utility of experimental and statistical approaches in determining the KIEs of rate-limiting steps in metabolic pathways. A similar framework could be used in the future to calculate such KIEs in additional methanogenic pathways, such as growth on methylamines, methyl sulfides, methoxylated compounds, and acetate to gain a better understanding of the parameters that control the isotopic composition of microbial methane in the environment. The approach can also be extended to other metabolisms and isotopic systems in which the rate limiting steps are known.

## Materials and Methods

### Strains and growth media

The *Methanosarcina acetivorans* strains used in this study were described previously and are listed in Table 1. We used WWM1 (Δ*hpt*) (54) as the “wildtype” *M. acetivorans* strain for all growth experiments. The strains WWM170, WWM176, and WWM184 express a synthetic operon comprised of *mtaC2B2A1, mtaC1B1A1*, and *mtaC3B3A1*, respectively, in a Δ*mtaC1B1A1*Δ*mtaC2B2A1*Δ*mtaC3B3A1*Δ*mtaA1*Δ*mtaA2* deletion mutant and under a tetracycline-inducible *mcr* promoter [Δ*hpt*::P*mcrB*-*tetR*-*ϕ*C31intattP::P*mcrB*(*tetO1*)-*mtaCXBXA1*, where ‘X’ stands for the 1, 2, or 3 isoforms (35)]. These strains, in principle, should only grow on methanol if the growth medium is supplemented with tetracycline to induce expression of the *mtaCXBXA1* operon. However, our growth experiments indicated that WWM170, WWM176, and WWM184 can grow on methanol even in the absence of tetracycline (Fig. S5). In addition, there was no observable difference in the growth rates when the mutants grown on 1, 10, and 100 μg/mL tetracycline (Fig. S2). Whole genome sequencing of WWM170, WWM176, and WWM184 (Tables S8-S10) revealed that each of them has a disruption in the *tetR* open reading frame, which leads to constitutive expression of *mtaCXBXA1* operon. Thus, the different tetracycline treatments were regarded as biological replicates.

All strains were grown in single-cell morphology at 37°C without shaking (HeraTherm General Protocol Microbiological Incubator, Thermo Fisher Scientific) in PIPES-buffered high-salt (HS) liquid medium (55) with a headspace of helium at an overpressure of 70 kPa. The medium was supplemented with 125 mM methanol as the sole carbon source. Anaerobic, sterile fresh stocks of tetracycline hydrochloride were prepared in deionized water before use and added to final concentrations between 1 to 100 μg/mL^−1^. Growth was conducted in 26 mL Balch tubes with 10 mL medium.

### Cultivation for isotopic assays

All growth experiments were initiated with stationary phase cultures pre-grown in 50 mM trimethylamine. The cells were transferred to HS medium with 125 mM methanol. Each experiment was initiated with 5 tubes for each replicate (3 replicates for each mutant strain) or with a total of 10 tubes for the WT. Growth was monitored with optical density measurements at 600 nm (OD_600_), using a UV-Vis Spectrophotometer (Genesys 50, Thermo Fisher Scientific) with a blank containing 10 mL uninoculated HS media (Fig. S2). Growth was arrested along the growth curve by injection of 1 mL 1 M NaOH solution to generate 5 representative time points from each replicate. Following NaOH injection, tubes were maintained at room temperature. The experiments were concluded when the OD_600_ of the cultures did not change for at least 48 hours. Tubes that contained tetracycline were wrapped in tin foil to avoid light-dependent degradation of tetracycline over time.

### DNA extraction and sequencing

We extracted the genomic DNA for whole genome re-sequencing of strains WWM170, WWM176, and WWM184 from 10 mL of stationary phase culture, using the Qiagen blood and tissue kit (Qiagen, Hilden, Germany). Genomic DNA sequencing was performed by SeqCenter (Pittsburgh, PA) where our samples were further processed using the Illumina DNA Prep kit. We mapped the sequencing reads (150 bp paired end reads) to a genome of the *M. acetivorans* strain WWM75 (39), using breseq v.0.35.5 (56). The complete list of the mutations that the breseq analysis identified is given in Tables S8-S10.

### Methane concentration determinations

CH_4_ and He concentrations in the headspace were measured using a Clarus 590 Gas Chromatograph (GC) (Perkin-Elmer) outfitted with a HayeSep 1/8’’ N60/80-6ft column (PerkinElmer) and a 100 μL sample loop, using Ar with a flow rate of 27 mL min^−1^ as a carrier gas. CH_4_ and He were detected with a flame ionization detector (FID) and thermal conductivity detector (TCD), respectively. Relative changes in CH_4_ content in the headspace during a given growth experiment was tracked as the ratio of the peak sizes of CH_4_ relative to He (Table S2).

### Measurement of carbon and hydrogen isotopic composition of methane

The methane carbon and hydrogen isotopic compositions (δ^13^C and δD) were determined at the Center for Isotope Geochemistry at the Lawrence Berkeley National Laboratory using a gas-chromatograph isotope ratio mass spectrometer (GC-IRMS) system (Thermo Scientific GC TraceGas Ultra system connected to a Thermo Scientific Delta V Plus IRMS), as described previously (50). Briefly, the gases were sampled using a gas-tight syringe and injected into a stainless-steel loop with a chosen size based on sample concentration (5 μL to 100 μL) attached on a 6-port valve (VICI-Valco). The methane was separated chromatographically on an HP-mole sieve fused silica capillary column (30 m X 0.32 mm) using He as carrier gas. Following chromatographic separation, CH_4_ was passed through a ceramic tube at 1200°C, combusted, and converted to CO_2_ for carbon isotopic measurements, or through a ceramic tube at 1420°C, pyrolyzed, and converted to H_2_ for hydrogen isotopic measurements. For hydrogen isotopic measurements, ceramic tubes were pre-conditioned by injecting 250 μL of pure methane three times the day before the measurement session. Measured δ^13^C and δD values were corrected for instrument linearity and external natural gas standards obtained from the United States Geological Service HCG-1, HCG-2, and HCG-3 (57) (Table S2). The δ^13^C and δD values of the methanol stocks were the same as those used in Gropp et al., 2025 (9) and measured as part of that study following methods in Lloyd et al., 2020 (58). Briefly, pure methanol was derivatized to CH_3_I using HI acid and then converted to chloromethane (CH_3_Cl) using excess silver chloride (AgCl). The isotopic composition of CH_3_Cl was measured using a MAT 253 Ultra (Thermo Scientific) at UC Berkeley and yielded a δD value of −49.0 ± 0.2‰ (1SE) and a δ^13^C value of −41.67 ± 0.01‰ (1SE) on the Vienna Standard Mean Ocean Water scale (VSMOW) and Vienna Peedee Belemnite scale (VPDB), respectively. The δD of the water (δD_H2O_) was measured for representative samples at the Center for Stable Isotope Biogeochemistry at UC Berkeley, via conversion of water to H_2_ using a hot chromium reactor unit (Thermo H/Device) connected to a Thermo Delta V Plus isotope ratio mass spectrometer. External water standards of known isotopic composition GFLES-2 (+159.9‰), GFLES-3 (+280.2‰), and GFLES-4 (+399.8‰) (USGS Reston) were used to correct samples to the VSMOW scale.

### In silico isotope enabled metabolic modeling of methylotrophic methanogenesis

To calculate the enzyme-specific KIEs, we used a reaction network of the methylotrophic pathway during growth on methanol that includes the minimal set of reactions needed to model the measured isotope effects that we described in the Results section (Fig. S1). The model includes a biomass sink from CH_3_-H_4_MPT, which we set at 10% of the total flux of methanol into the pathway based on prior work (12). Together with the 3:1 ratio of the products CH_4_:CO_2_, this sink determines the disproportionation ratio of CH_3_-S-CoM between MCR and MTR. As mentioned in the Results section, we allowed partial reversibility in the oxidative branch following previously reported values of ∼0.05 net reversibility (9). The reversibility of the *i*th reaction ‘*r*_*i*_’is defined here as the ratio of the backward to forward fluxes of the reaction (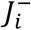 and 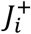, respectively), where 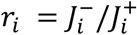. We note that this value represents the net reversibility for hydrogen isotopes, which can be calculated as the average of the reversibility of each step in the pathway: 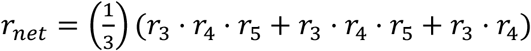, where the numbers correspond to the reactions in Fig. S1. Since we assumed here that the reaction catalyzed by methylene– tetrahydromethanopterin dehydrogenase (MTD) is irreversible, i.e., *r*_5_ = 0, MTR and MER were assigned with a reversibility of 0.4 to obtain a net reversibility of 0.05. We provide the results of a sensitivity analysis in regard to the degree of reversibility in Table S6, and the isotopic model parameters in Table S4.

### Calculation of kinetic isotopic effects (KIEs)

We estimated ^13^ε and ^2^ε values using the MCMC algorithm implemented in the *emcee* package in Python (59). To determine which of these parameters could be resolved with the MCMC algorithm given the experimental results, we generated a synthetic dataset with the model described in the ‘In silico isotope enabled metabolic modeling of methylotrophic methanogenesis’ section. We found that the model reproduces the ^13^ε_MTA_ and ^2^ε_MTA_ and the ratios of the carbon KIEs of MCR and MTR (^13^KIE_MCR/MTR_) of the synthetic data set. The algorithm works as follows: Using predefined boundaries for the KIEs and the initial δ^13^C_CH3OH_, δD_CH3OH_, and δD_H2O_ values, the MCMC algorithm solves the isotopic model multiple times while minimizing the distance from the modeled vs. measured δ^13^C_CH4_ and δD_CH4_ values. The MCMC parameters in our simulations were 100 walkers, 600 steps, and 400 burn-in discarded steps resulting in overall 20,000 effective samples. The posterior distributions of the KIEs were extracted from the MCMC algorithm and were used to estimate the uncertainty of the KIEs which were defined by the 0.025 and 0.975 percentiles (95% of the posterior model results). In experiments involving mutant strains WWM170, WWM176, and WWM184, three replicates were conducted. For these, we pooled the posterior distributions to estimate uncertainties. Statistical significance was assessed by calculating the element-wise difference between parameter posterior distributions. A difference was considered significant if the 95% credible interval of the difference distribution excluded zero. When estimating the KIEs using MCMC we incorporate the uncertainty in the measured product yield (*f*) and of the isotopic values, by introducing noise that is proportional to the standard deviation of those parameters. Specifically, we use a relative uncertainty of 2% (1SD) in our *f* values, 0.07‰ in our δ^13^C_CH4_ values (1SD), and 2‰ for our δD_CH4_ values (1SD). In our comparisons of the synthetic datasets and previously published work we use similar uncertainties to be internally consistent.

## Acknowledgments

We thank Prof. William Metcalf from the University of Illinois Urbana-Champaign for providing us with the strains that were used in this study, Frank Cai for helping to analyze the headspace CH_4_ and He concentrations, and Dr. Markus Bill from the Lawrence Berkeley National Lab for helping in analyzing the carbon and hydrogen isotopic compositions. JG acknowledges support from the European Molecular Biology Organization (EMBO) and from the Institute for Environmental Sustainability (IES). DAS acknowledges support from the Alfred P. Sloan Research Fellowship sponsored by the Sloan Foundation. DDN acknowledges funding from the Searle Scholars Program sponsored by the Kinship Foundation, the Rose Hills Innovator Grant, the Beckman Young Investigator Award sponsored by the Arnold and Mabel Beckman Foundation, the Alfred P. Sloan Research Fellowship sponsored by the Sloan Foundation, the Simons Foundation Early Career Investigator in Marine Microbial Ecology and Evolution Award, the Packard Fellowship in Science and Engineering sponsored by the David and Lucille Packard Foundation, and the Department of Energy through project number S589706. DDN is a Chan-Zuckerberg Biohub – San Francisco Investigator. The funders had no role in the conceptualization and writing of this manuscript or the decision to submit the work for publication.

## Conflicts of Interest

The authors declare no conflicts of interest.

## Author contributions

- Conceptualization, Writing – review and editing, Funding acquisition: JG, DAS, DDN
- Writing – original draft: JG
- Investigation, Software: JG
- Supervision: DAS, DDN

## Data and materials availability

All data are available in the manuscript or the supplementary materials. The DNA sequences of the mutant strains were deposited in the NCBI database at https://www.ncbi.nlm.nih.gov/bioproject/1444202.

## Supplementary text

### An isotopic mass balance of methylotrophic methanogenesis on methanol

The following equations were used to track the time derivatives of the [CH_3_OH], [CH_4_], and [CO_2_], and their ^13^C/^12^C and D/H ratios. The reactions’ numbers correspond to the reactions’ network as presented in Fig. S1.

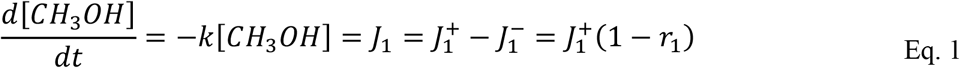

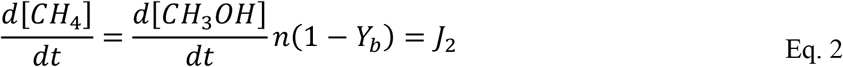

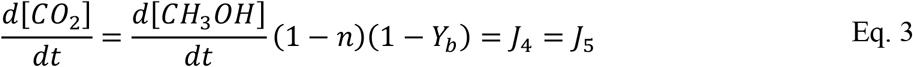

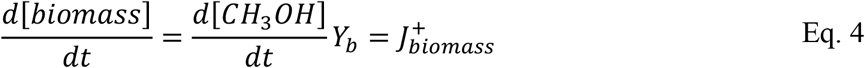

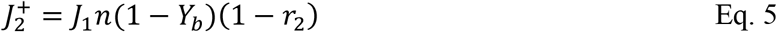

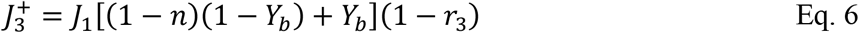

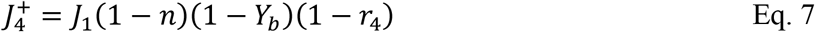

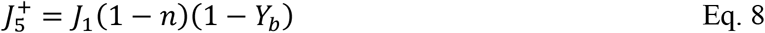

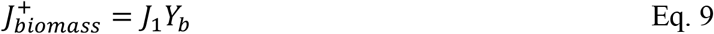

where *J*_*i*_ is the metabolic flux of the *i*th reaction, and 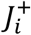 and 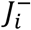 denote its forward and reverse fluxes, respectively, *Y*_*b*_ is the biomass sink, and *n* is the fraction of CH_3_-S-CoM molecules reduced to CH_4_ relative to CO_2_. The reversibility of the *i*th reaction *r*_*i*_ is defined by 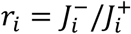. In this work we used *n* = 0.75 which represents a CH_4_:CO_2_ ratio of 3:1, and *Y*_*b*_ = 0.1 (12).

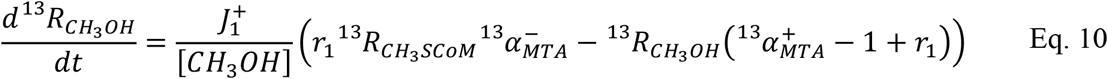

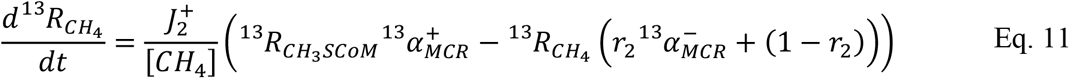

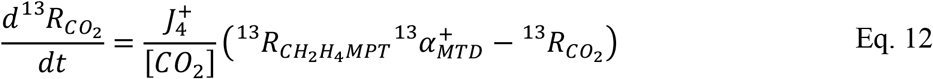

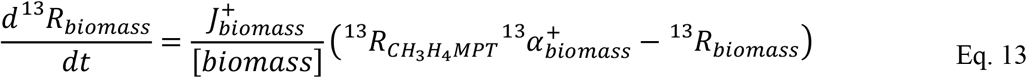

Here ^13^*R*_*j*_ is the ^13^C/^12^C ratio of the *j*th metabolite, and 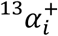 and 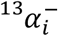 are the forward and reverse carbon KIEs of the i*th* reaction, respectively.

We assumed that the intermediate metabolites are at a steady state between the integration steps so that:

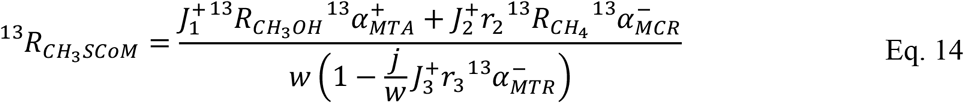

where:

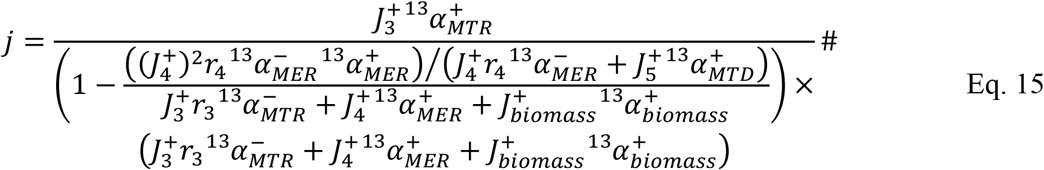

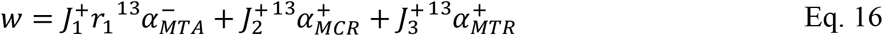

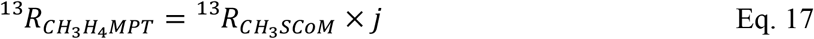

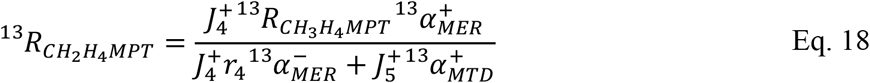

We constructed a similar mass balance for hydrogen isotopes. Here the KIEs are denoted by the ‘s’ and ‘p’ superscripts that represent secondary and primary hydrogen KIEs, respectively.

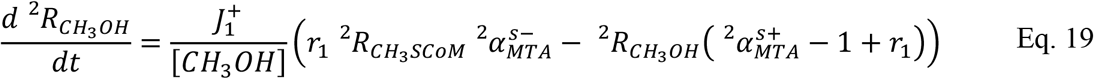

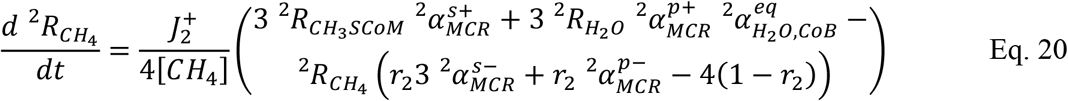

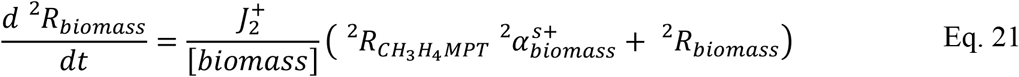

Similarly to the carbon isotope mass balance, we assume that the intermediate metabolites are at a steady state between the integration steps so that:

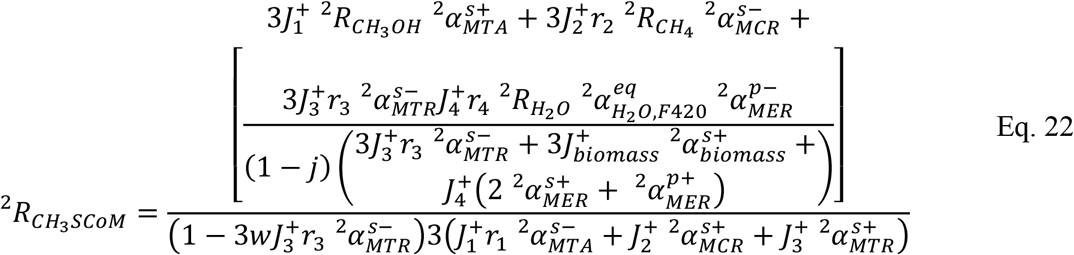

where:

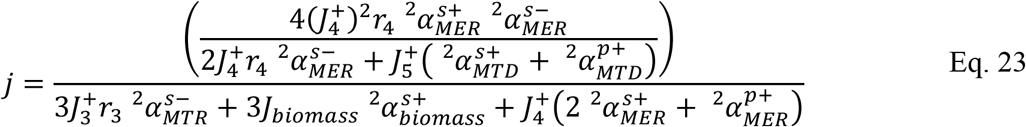

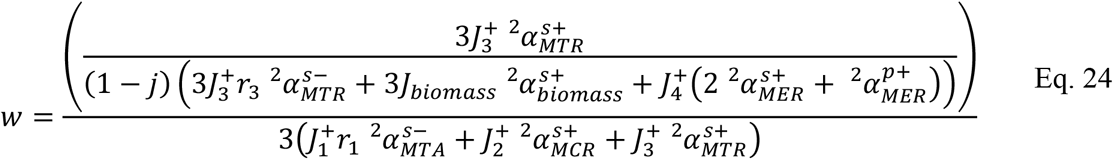

## Supplementary Figures

**Fig. S1.**
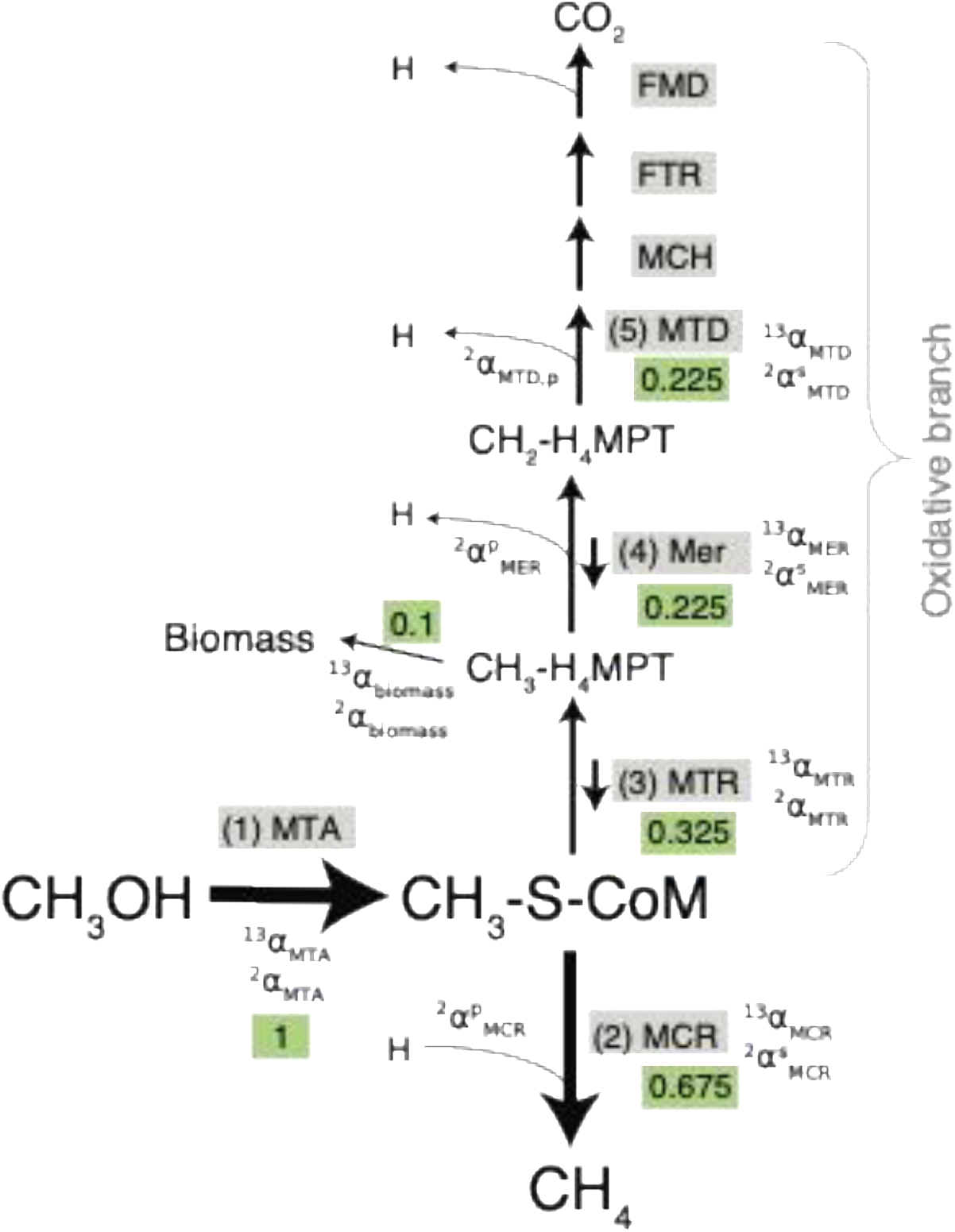
Metabolic network for the hydrogen and carbon isotopic mass balance. The numbers and notations for the KIEs correspond to those used in Eqs. S1–S23. The green boxes represent the relative flux of carbon in the pathway. Reactions that break a C-H bond have both secondary and primary KIEs, denoted by ‘s’ and ‘p’ next to the α, respectively. The model assumes a 3:1 ratio of CH_4_:CO_2_ production, a biomass sink that is 10% of the total flux of methanol, and partial reversibility in the oxidative branch in the reactions catalyzed by MTR and Mer. The reactions catalyzed by MCH, FTR, and FMD do not affect the isotopic composition in methane because we assume that MTD is irreversible and thus they do not control the isotopic composition of the final methane. Abbreviations for enzyme complexes names and metabolites: CH_3_-S-CoM – methyl coenzyme M; H_4_MPT – tetrahydromethanopterin; MTA – methanol specific methyltransferase; MCR – methyl-CoM reductase; MTR – N^5^-methyl-H_4_MPT:coenzyme M methyltransferase; Mer – N^5^,N^10^-methylene-H_4_MPT:coenzyme-F420 reductase; MTD – N^5^,N^10^-methylene-H_4_MPT:coenzyme-F420 reductase. MCH – methenyl-H_4_MPT cyclohydrolase; FTR – formylmethanofuran:H_4_MPT formyltransferase; FMD – formylmethanofuran dehydrogenase.

**Fig. S2.**
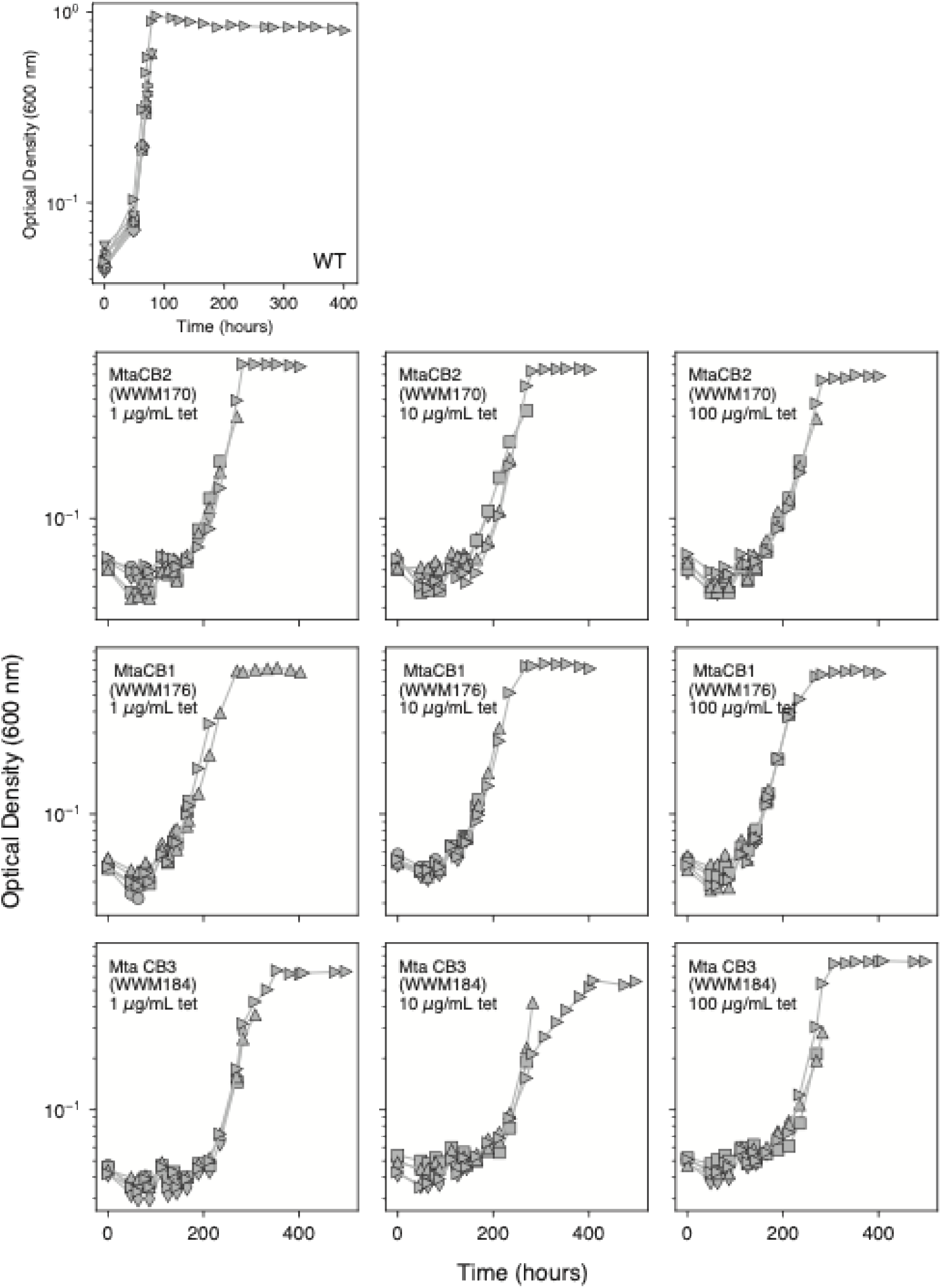
Growth curves of WT and mutant strains. Optical density (OD) measurements were taken at 600 nm. The different symbols represent the replicates, where each symbol represents a single tube, with a total of 10 tubes for the WT and 5 per each mutant strain on a given tetracycline (tet) concentration. Tubes were sacrificed during the experiment by injection of 1 mL NaOH 1M for headspace analysis.

**Fig. S3.**
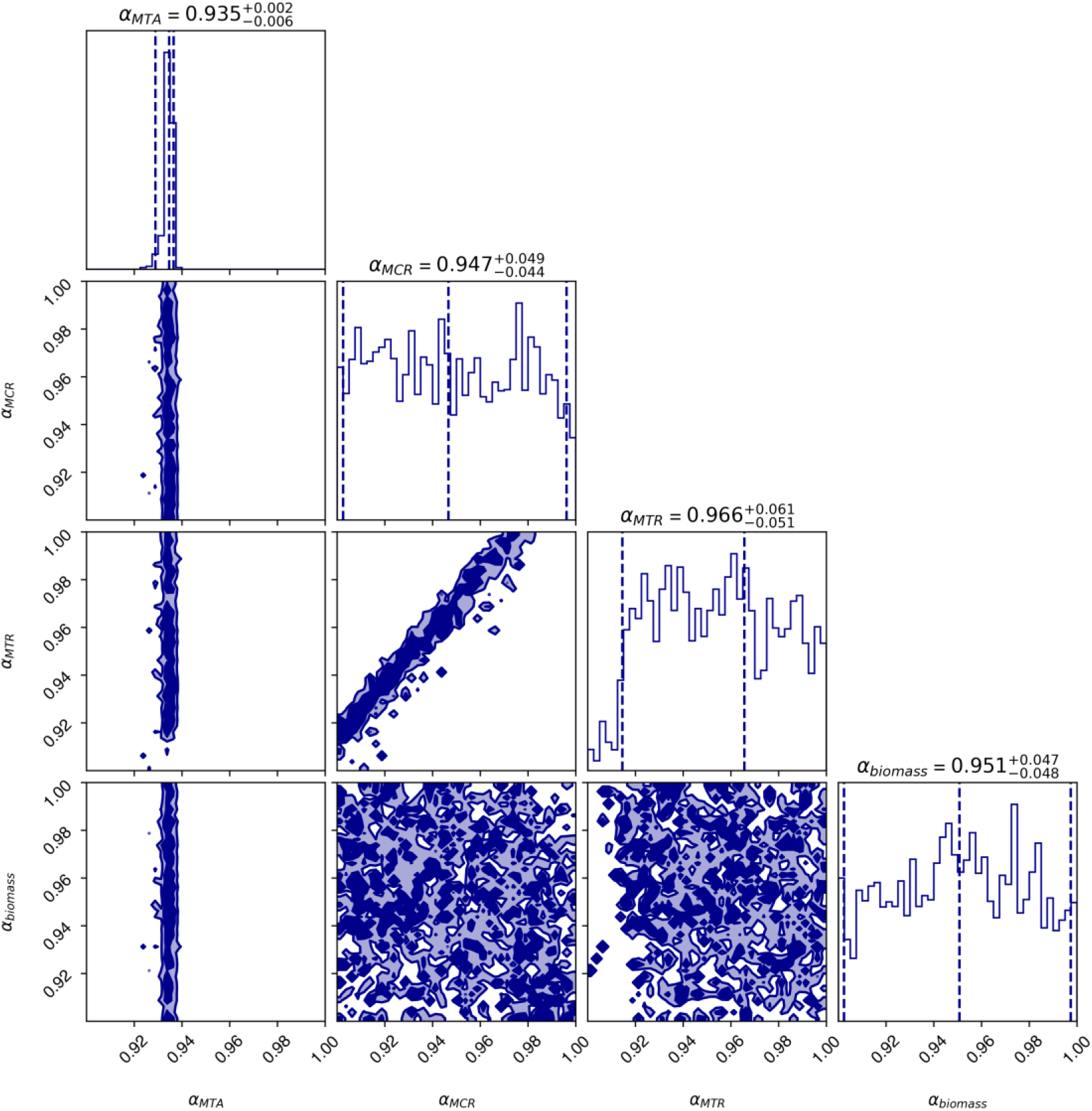
Test of carbon KIEs fits using the MCMC algorithm with synthetic data. Here, the KIEs are denoted as ‘α’.The predefined KIEs were ^13^*α*_*MTA*_ = 0.935, ^13^*α*_*MCR*_ = 0.960, ^13^*α*_*MTR*_ = 0.980 (i.e., ^13^*α*_*MCR*_/^13^*α*_*MTR*_ = 0.980), and ^13^*α*_*biomass*_ of 0.98. The posterior distributions of 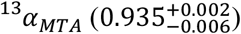 match the predefined value, while ^13^*α*_*MCR*_ and ^13^*α*_*MTR*_ do not match the predefined values and yield much larger uncertainties. However, the ratios of ^13^*α*_*MCR*_ and ^13^*α*_*MTR*_ yield a value of 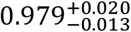, which matches the predefined value of 0.980. Analysis was performed using 250 walkers for 400 steps. The first 200 steps were discarded as burn-in, resulting in 50,000 effective samples used for parameter estimation. The diagonal histograms show the one-dimensional normalized posterior distributions for each parameter. The off-diagonal panels display the two-dimensional posterior distributions for all pairs of parameters, illustrating their correlations. Contours represent 68% (dark blue) and 95% (light blue) credible regions.

**Fig. S4.**
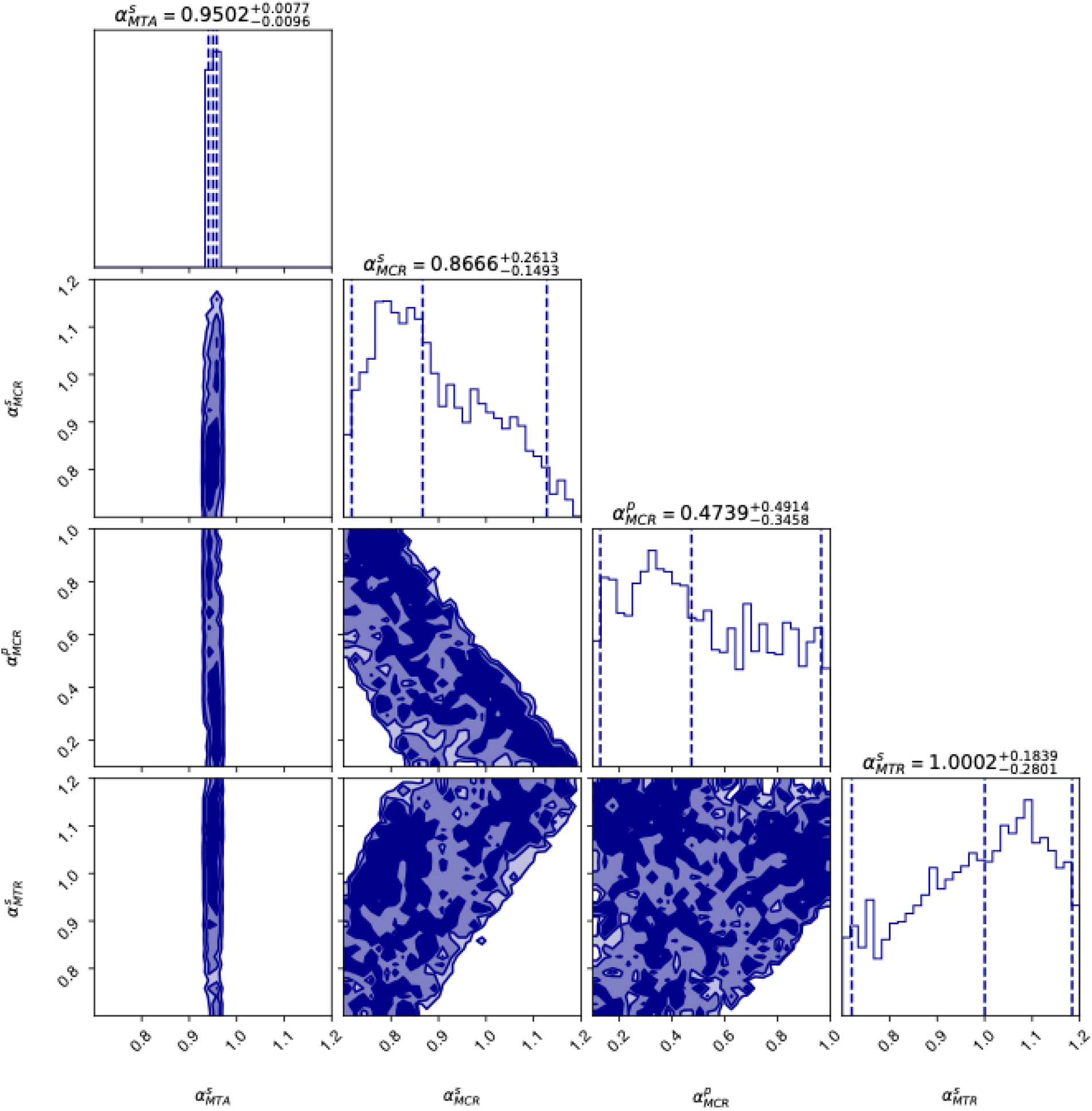
Test of hydrogen KIEs using the MCMC algorithm with synthetic data. Here, the KIEs are denoted as ‘α’. The predefined KIEs were 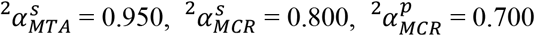, and 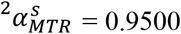. Analysis was performed using 10 walkers for 10,000 steps. The first 200 steps were discarded as burn-in, resulting in 98,000 effective samples used for parameter estimation. The posterior distributions of 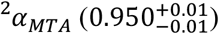 match the predefined value. The three additional KIEs yield posterior distribution with large error estimates, and values that do not match the predefined values. The diagonal histograms show the one-dimensional normalized posterior distributions for each parameter. The off-diagonal panels display the two-dimensional posterior distributions for all pairs of parameters, illustrating their correlations. Contours represent 68% (dark blue) and 95% (light blue) credible regions. Note that the scales are different between the parameters, representing the prescribed bounds of each parameter.

**Fig. S5.**
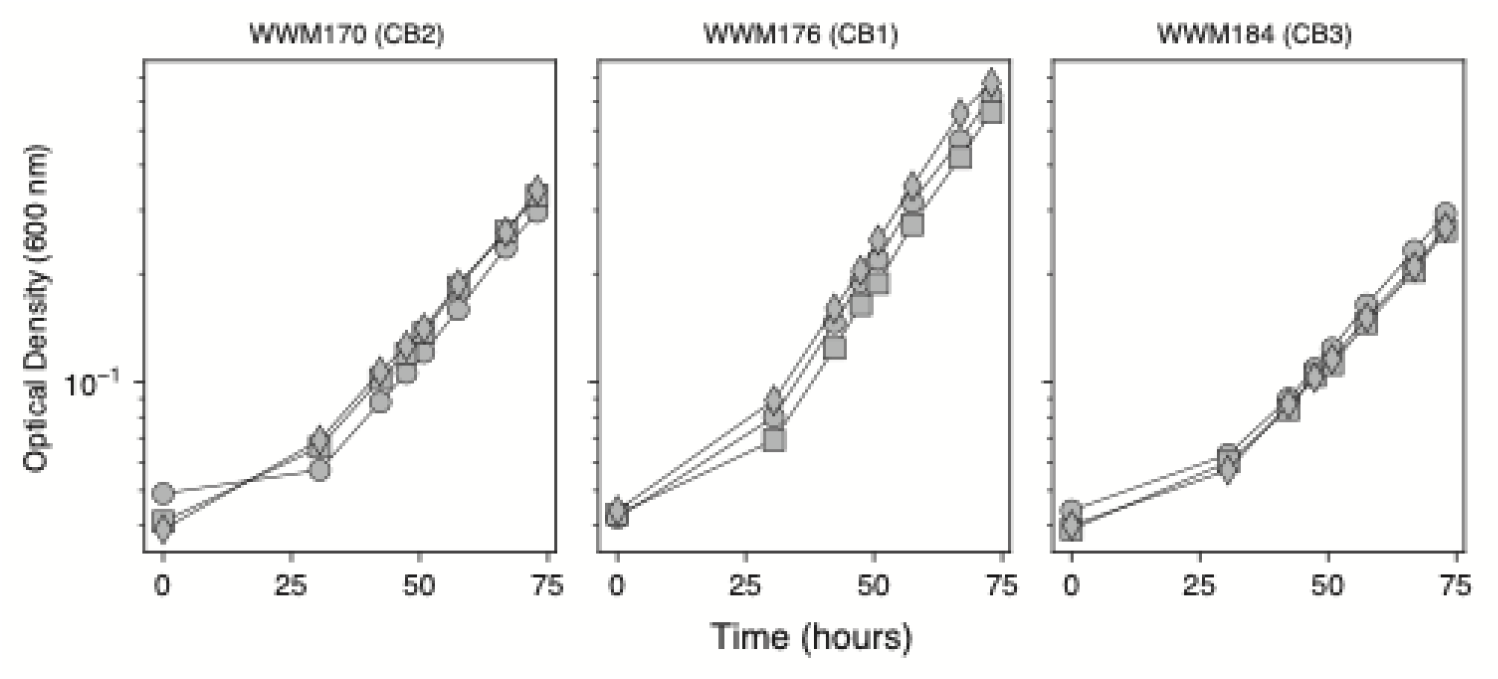
Mutant strains grow exponentially without tetracycline addition. Optical density (OD) measurements were taken at 600 nm. Strains were thawed from freezer stocks to HS-TMA media (50 mM) and then acclimated for growth on HS media with 60 mM methanol for one generation. Then 0.2 mL of late exponential phase culture was transferred in triplicates to HS media with 60 mM methanol but no tetracycline (replicates represented by the symbols).

## List of Supplementary Tables

Table S1. Data compilation from previous work used to plot Fig. 1A.

Table S2. Growth data, carbon and hydrogen isotopic compositions of methane, and product yields.

Table S3. Growth parameters of the different strains used in this study.

Table S4. Isotopic parameters for the model.

Table S5. Fitting δ^13^C_CH3OH_ and δ^13^C_CH4_ from Ref. (13).

Table S6. Sensitivity analysis for reversibility and the size of the biomass sink.

Table S7. The contribution of the KIE of MTA to the overall isotopic offset.

Table S8. Whole genome sequence analysis with Breseq for WWM170.

Table S9. Whole genome sequence analysis with Breseq for WWM176.

Table S10. Whole genome sequence analysis with Breseq for WWM184.

## References

1. Whiticar MJ. 1999. Carbon and hydrogen isotope systematics of bacterial formation and oxidation of methane. Chemical Geology 161:291–314.

2. Douglas PMJ, Stratigopoulos E, Park S, Phan D. 2021. Geographic variability in freshwater methane hydrogen isotope ratios and its implications for global isotopic source signatures. Biogeosciences 18:3505–3527.

3. Michel SE, Lan X, Miller J, Tans P, Clark JR, Schaefer H, Sperlich P, Brailsford G, Morimoto S, Moossen H, Li J. 2024. Rapid shift in methane carbon isotopes suggests microbial emissions drove record high atmospheric methane growth in 2020–2022. Proc Natl Acad Sci USA 121:e2411212121.

4. Krzycki JA, Kenealy WR, Deniro MJ, Zeikus JG. 1987. Stable Carbon Isotope Fractionation by Methanosarcina barkeri during Methanogenesis from Acetate, Methanol, or Carbon Dioxide-Hydrogen. Applied and environmental microbiology 53:2597–9.

5. Hayes JM. 2001. Fractionation of Carbon and Hydrogen Isotopes in Biosynthetic Processes. Reviews in Mineralogy and Geochemistry 43:225–277.

6. Valentine DL, Chidthaisong A, Rice A, Reeburgh WS, Tyler SC. 2004. Carbon and hydrogen isotope fractionation by moderately thermophilic methanogens. Geochimica et Cosmochimica Acta 68:1571–1590.

7. Gropp J, Jin Q, Halevy I. 2022. Controls on the isotopic composition of microbial methane. Science Advances 8:eabm5713.

8. Mayumi D, Tamaki H, Kato S, Igarashi K, Lalk E, Nishikawa Y, Minagawa H, Sato T, Ono S, Kamagata Y, Sakata S. 2024. Hydrogenotrophic methanogens overwrite isotope signals of subsurface methane. Science 386:1372–1376.

9. Gropp J, Bill M, Lloyd MK, Stein R, Nayak DD, Stolper DA. 2025. Modulation of methyl-coenzyme M reductase expression alters the isotopic composition of microbial methane. Science 10.1126/science.adu2098.

10. Miller HM, Chaudhry N, Conrad ME, Bill M, Kopf SH, Templeton AS. 2018. Large carbon isotope variability during methanogenesis under alkaline conditions. Geochimica et Cosmochimica Acta 237:18–31.

11. Rosenfeld WD, Silverman SR. 1959. Carbon Isotope Fractionation in Bacterial Production of Methane. Science 130:1658–1659.

12. Londry KL, Dawson KG, Grover HD, Summons RE, Bradley AS. 2008. Stable carbon isotope fractionation between substrates and products of Methanosarcina barkeri. Organic Geochemistry 39:608–621.

13. Penger J, Conrad R, Blaser M. 2012. Stable carbon isotope fractionation by methylotrophic methanogenic archaea. Applied and environmental microbiology 78:7596–602.

14. Games LM, Hayes JM, Gunsalus RP. 1978. Methane-producing bacteria: natural fractionations of the stable carbon isotopes. Geochimica et Cosmochimica Acta 42:1295–1297.

15. Fuchs G, Thauer R, Ziegler H, Stichler W. 1979. Carbon isotope fractionation by Methanobacterium thermoautotrophicum. Archives of Microbiology 120:135–139.

16. Belyaev SS, Wolkin R, Kenealy WR, Deniro MJ, Epstein S, Zeikus JG. 1983. Methanogenic bacteria from the bondyuzhskoe oil field: general characterization and analysis of stable-carbon isotopic fractionation. Applied and environmental microbiology 45:691–7.

17. Balabane M, Galimov E, Hermann M, Létolle R. 1987. Hydrogen and carbon isotope fractionation during experimental production of bacterial methane. Organic Geochemistry 11:115–119.

18. Gelwicks JT, Risatti JB, Hayes JM. 1994. Carbon isotope effects associated with aceticlastic methanogenesis. Applied and environmental microbiology 60:467–72.

19. Botz R, Pokojski HD, Schmitt M, Thomm M. 1996. Carbon isotope fractionation during bacterial methanogenesis by CO2 reduction. Organic Geochemistry 25:255–262.

20. Summons RE, Franzmann PD, Nichols PD. 1998. Carbon isotopic fractionation associated with methylotrophic methanogenesis. Organic Geochemistry 28:465–475.

21. House CH, Schopf JW, Stetter KO. 2003. Carbon isotopic fractionation by Archaeans and other thermophilic prokaryotes. Organic Geochemistry 34:345–356.

22. Penning H, Claus P, Casper P, Conrad R. 2006. Carbon isotope fractionation during acetoclastic methanogenesis by Methanosaeta concilii in culture and a lake sediment. Applied and Environmental Microbiology 72:5648–52.

23. Takai K, Nakamura K, Toki T, Tsunogai U, Miyazaki M, Miyazaki J, Hirayama H, Nakagawa S, Nunoura T, Horikoshi K. 2008. Cell proliferation at 122 degrees C and isotopically heavy CH4 production by a hyperthermophilic methanogen under high-pressure cultivation. Proceedings of the National Academy of Sciences of the United States of America 105:10949–54.

24. Goevert D, Conrad R. 2009. Effect of substrate concentration on carbon isotope fractionation during acetoclastic methanogenesis by Methanosarcina barkeri and M. acetivorans and in rice field soil. Applied and Environmental Microbiology 75:2605–2612.

25. Penger J, Conrad R, Blaser M. 2014. Stable carbon isotope fractionation of six strongly fractionating microorganisms is not affected by growth temperature under laboratory conditions. Geochimica et Cosmochimica Acta 140:95–105.

26. Li J, Ash JL, Cobban A, Kubik BC, Rizzo G, Thompson M, Guibourdenche L, Berger S, Morra K, Lin Y, Mueller EP, Masterson AL, Stein R, Fogel M, Torres MA, Feng X, Holden JF, Martini A, Welte CU, M. Jetten MS, Young ED, Leavitt WD. 2025. The Clumped Isotope Signatures of Multiple Methanogenic Metabolisms. Environ Sci Technol 59:13798–13810.

27. Zhuang G-C, Elling FJ, Nigro LM, Samarkin V, Joye SB, Teske A, Hinrichs K-U. 2016. Multiple evidence for methylotrophic methanogenesis as the dominant methanogenic pathway in hypersaline sediments from the Orca Basin, Gulf of Mexico. Geochimica et Cosmochimica Acta 187:1–20.

28. Xiao K-Q, Beulig F, Røy H, Jørgensen BB, Risgaard-Petersen N. 2018. Methylotrophic methanogenesis fuels cryptic methane cycling in marine surface sediment. Limnology and Oceanography 63:1519–1527.

29. Narrowe AB, Borton MA, Hoyt DW, Smith GJ, Daly RA, Angle JC, Eder EK, Wong AR, Wolfe RA, Pappas A, Bohrer G, Miller CS, Wrighton KC. 2019. Uncovering the Diversity and Activity of Methylotrophic Methanogens in Freshwater Wetland Soils. mSystems 4:e00320–19.

30. Hanna E, Keller JK, Chang D, de Bruyn W, Zalman C. 2020. The potential importance of methylated substrates in methane production within three northern Minnesota peatlands. Soil Biology and Biochemistry 150:107957.

31. Schorn S, Ahmerkamp S, Bullock E, Weber M, Lott C, Liebeke M, Lavik G, Kuypers MMM, Graf JS, Milucka J. 2022. Diverse methylotrophic methanogenic archaea cause high methane emissions from seagrass meadows. PNAS 119.

32. Zhuang G, Liu Q, Yuan L, Wu B, Sui X, Zhou Z, Wang J, Dong L, Liu X, Yang G, Wang F. 2025. Controls and contributions of methylotrophic methanogenesis to methane production in coastal sediments. Limnology & Oceanography lno.70269.

33. Krause SJE, Wipfler R, Liu J, Yousavich DJ, Robinson D, Hoyt DW, Orphan VJ, Treude T. 2025. Spatial evidence of cryptic methane cycling and methylotrophic metabolisms along a land–ocean transect in salt marsh sediment. Geochimica et Cosmochimica Acta 404:53–71.

34. Krause SJE, Treude T. 2021. Deciphering cryptic methane cycling: Coupling of methylotrophic methanogenesis and anaerobic oxidation of methane in hypersaline coastal wetland sediment. Geochimica et Cosmochimica Acta 10.1016/j.gca.2021.03.021.

35. Opulencia RB, Bose A, Metcalf WW. 2009. Physiology and Posttranscriptional Regulation of Methanol:Coenzyme M Methyltransferase Isozymes in Methanosarcina acetivorans C2A. Journal of Bacteriology 191:6928–6935.

36. Bose A, Pritchett MA, Rother M, Metcalf WW. 2006. Differential Regulation of the Three Methanol Methyltransferase Isozymes in Methanosarcina acetivorans C2A. Journal of Bacteriology 188:7274–7283.

37. Scheller S, Goenrich M, Thauer RK, Jaun B. 2013. Methyl-coenzyme M reductase from methanogenic archaea: isotope effects on the formation and anaerobic oxidation of methane. Journal of the American Chemical Society 135:14975–84.

38. Jin Q, Wu Q, Shapiro BM, McKernan SE. 2022. Limited Mechanistic Link Between the Monod Equation and Methanogen Growth: a Perspective from Metabolic Modeling. Microbiology Spectrum 10.1128/spectrum.02259-21.

39. Guss AM, Rother M, Zhang JK, Kulkkarni G, Metcalf WW. 2008. New methods for tightly regulated gene expression and highly efficient chromosomal integration of cloned genes for Methanosarcina species. Archaea 2:193–203.

40. Sivan M, Wallenius AJ, Röckmann T, Jetten MSM, Slomp CP, Greule M, Keppler F, Gilbert A, Yamada K, Moonen RPJ, Popa ME. 2025. Pathway-specific bulk and clumped isotope signatures of methane production in marine sediment incubations. Biogeochemistry 168:90.

41. Bowen GJ, Ehleringer JR, Chesson LA, Stange E, Cerling TE. 2007. Stable isotope ratios of tap water in the contiguous United States. Water Resources Research 43:2006WR005186.

42. Wegener G, Gropp J, Taubner H, Halevy I, Elvert M. 2021. Sulfate-dependent reversibility of intra-cellular reactions explains the opposing isotope effects in the anaerobic oxidation of methane. Science Advances 7:eabe4939.

43. Klinman JP. 1978. Kinetic Isotope Effects in Enzymology, p. 415–494. In Meister, A (ed.), Advances in Enzymology - and Related Areas of Molecular Biology, 1st ed. Wiley.

44. Cleland WW. 2005. The use of isotope effects to determine enzyme mechanisms. Archives of Biochemistry and Biophysics 433:2–12.

45. Hagemeier CH, Krër M, Thauer RK, Warkentin E, Ermler U. 2006. Insight into the mechanism of biological methanol activation based on the crystal structure of the methanol-cobalamin methyltransferase complex. Proc Natl Acad Sci USA 103:18917–18922.

46. Westaway KC. 2006. Using kinetic isotope effects to determine the structure of the transition states of SN2 reactions, p. 217–273. In Advances in Physical Organic Chemistry. Elsevier.

47. Gray CH, Coward JK, Schowen KB, Schowen RL. 1979. .alpha.-Deuterium and carbon-13 isotope effects for a simple, intermolecular sulfur-to-oxygen methyl-transfer reaction. Transition-state structures and isotope effects in transmethylation and transalkylation. J Am Chem Soc 101:4351–4358.

48. Hegazi MF, Borchardt RT, Schowen RL. 1979. .alpha.-Deuterium and carbon-13 isotope effects for methyl transfer catalyzed by catechol O-methyltransferase. SN2-like transition state. J Am Chem Soc 101:4359–4365.

49. Linscott JA, Kapilashrami K, Wang Z, Senevirathne C, Bothwell IR, Blum G, Luo M. 2016. Kinetic isotope effects reveal early transition state of protein lysine methyltransferase SET8. Proc Natl Acad Sci USA 113.

50. Turner AC, Korol R, Eldridge DL, Bill M, Conrad ME, Miller TF, Stolper DA. 2021. Experimental and theoretical determinations of hydrogen isotopic equilibrium in the system CH4-H2-H2O from 3 to 200°C. Geochimica et Cosmochimica Acta 1:1.

51. Ono S, Rhim JH, Ryberg EC. 2022. Rate limits and isotopologue fractionations for microbial methanogenesis examined with combined pathway protein cost and isotopologue flow network models. Geochimica et Cosmochimica Acta 10.1016/j.gca.2022.03.017.

52. Jautzy JJ, Douglas PMJ, Xie H, Eiler JM, Clark ID. 2021. CH4 isotopic ordering records ultra-slow hydrocarbon biodegradation in the deep subsurface. Earth and Planetary Science Letters 562:116841.

53. Gropp J, Iron MA, Halevy I. 2021. Theoretical estimates of equilibrium carbon and hydrogen isotope effects in microbial methane production and anaerobic oxidation of methane. Geochimica et Cosmochimica Acta 295:237–264.

54. Pritchett MA, Metcalf WW. 2005. Genetic, physiological and biochemical characterization of multiple methanol methyltransferase isozymes in Methanosarcina acetivorans C2A. Molecular Microbiology 56:1183–1194.

55. Sowers KR, Boone JE, Gunsalus RP. 1993. Disaggregation of Methanosarcina spp. and Growth as Single Cells at Elevated Osmolarity. Applied and Environmental Microbiology 59:3832–3839.

56. Deatherage DE, Barrick JE. 2014. Identification of Mutations in Laboratory-Evolved Microbes from Next-Generation Sequencing Data Using breseq, p. 165–188. In Sun, L, Shou, W (eds.), Engineering and Analyzing Multicellular Systems. Springer New York, New York, NY.

57. Dias RF, Coleman DD, Ellis GS. 2022. Stable Isotopic Data (Delta13C and Delta2H) for Reference Materials HCG-1, HCG-2, and HCG-3. [object Object].

58. Lloyd MK, Eldridge DL, Stolper DA. 2020. Clumped 13CH2D and 12CHD2 compositions of methyl groups from wood and synthetic monomers: methods, experimental and theoretical calibrations, and initial results. Geochimica et Cosmochimica Acta 10.1016/j.gca.2020.10.008.

59. Foreman-Mackey D, Hogg DW, Lang D, Goodman J. 2013. emcee: The MCMC hammer 125:306.

